# Differences in self-reported benefits for student-artist versus faculty experiences in a virtual artist-in-residence program

**DOI:** 10.1101/2022.09.14.507965

**Authors:** Skylar Cuevas, Qi (Kathy) Liu, Helen Qian, Max E. Joffe, Karisa Calvitti, Megan Schladt, Eric P. Skaar, Kendra H. Oliver

## Abstract

The value of science communication in engaging the public has been well established. While many new programs bridge the arts and sciences, conducting comprehensive examination of exiting art-science programs can produce more efficient training and program development guidance for improving visual communications in the sciences. Here, we recruited a variety of scientists and artists to collaborate in creating visual science communication products over three summers. Using survey data, we performed qualitative and quantitative analyses to define sources for negative and positive experiences and outcomes from the Vanderbilt Institute for Infection, Immunology, and Inflammation (VI4) Artist-in-Residence (AiR) program. Further, we analyze responses from participants, student-artists and faculty, to specify areas for improvement and areas successful in producing a positive experience and outcome in an AiR program. We found that time and virtual delivery of the program could be modified to improve the experience. Additionally, we found that student participants had more positive responses about “learning something new” from the program than faculty members. However, the most surprising aspect of our analysis suggests that for both “way of thinking” and “science communication to the public or general audience,” there may be more significant beneficial gains for faculty compared to students. We conclude this analysis with suggestions to enhance the benefits and outcomes of an AiR program and ways to minimize the difficulties, such as communication and collaboration, faced by participants and program designers.

## INTRODUCTIONS

Over the past few years, there has been a renaissance for science-based art programs and artist-in-residence (AiR) experiences[1–3]. The Vanderbilt Institute for Infection, Immunology, and Inflammation AiR program brings together people from multiple backgrounds and interests to produce visual science communication products. One of the main goals for the AiR program is to create an environment for people to collaborate and gain exposure to new communities and visual communication approaches. Based on previous publications, partnerships appear to be most valuable when scientists and artists have a shared stake in a project, facilitating their ability to jointly communicate, design, and critique the work[4,5]. Anecdotally, AiR experiences can prompt novel ways of thinking for artists and scientists alike and result in powerful visual science communication products. Although many of these programs aim to engage the general public in science, these experiences also impact and transform the participants, leading to innovative products that can turn science into captivating visuals[6].

Training in visual science communication is essential yet typically neglected within scientific training. This is exemplified by Estrada and Davis who identified that visual materials have typically been treated as an add-on instead of being an integrated aspect of science communication[7]. Visual science communication encompasses all images, graphics, and visual representations created to communicate an element of scientific knowledge. Guidance on improving the visual aspects of science communication range from step-by-step-style instructions to hyper-focused aspects of data visualization[8]. The value of science communication in engaging the public has been well established[9–12]; thus, having tools to increase the effectiveness of these efforts is imperative[13,14]. The public impact of science images is evident throughout the media, with innovative technologies playing a vital role in representing science. Visual literacy, defined here as a holistic construct encompassing visual thinking, learning, and communication, is a critical ingredient in the effective communication of science among expert and lay audiences[15]. However, within peer-reviewed literature, guidance on improving and assessing the effectiveness of written and oral presentation skills has dominated science communication research[16–20]. Less emphasis, however, has been placed on enhancing the visual aspects of science communication.

While peer-to-peer communications have long employed graphs and figures for journal publications and conference posters, the standard practice for academic and research institutions to hire graphic experts for assistance has largely fallen out of favor[21]. With the current high demand to connect with non-specialist audiences, more publications have been surfacing with guidance on improving various facets of visual communications, drawing specifically on incorporating graphic design principles. Many of these publications, however, are either limited in scope, are proved an excellent call to action with a lack of visuals (though it is a commentary), are aimed at one specific type of visual medium such as a poster presentation[22], or are hyper-focused on one specialized component such as producing imagery via understanding concepts in data visualization [8,21,23–25]. Considering these isolated approaches, conducting a comprehensive examination in exiting AiR programs can inform more efficient guidance on training and program development for improving visual communications in the sciences.

For art-science programs and AiR programs, there is a lack of information related to the program designs and benefits for artists and science faculty. At present, support for the educational benefits of art-science partnerships is anecdotal [6,26]. There have been surprisingly few attempts to test the widely held assumption that studying the arts makes one more creative in general [27]. And while the visual arts can develop students’ creativity, objectivity, perseverance, spatial reasoning, and observational acuity—key science skills—it is not clear whether skills developed through artistic pursuits can transfer to other fields[27–29]. Nevertheless, there are compelling reports of collaborations pre-undergraduate education and professional levels that have enriched general audience engagement and the scientists and artists at the center of the work[30,31]. These projects suggest that combining art and science can have transformative effects. We believe that the lingering question of knowledge transfer is all the more reason to develop projects where the boundaries between art and science are blurred, as merging two traditionally separate subjects can yield unexpected rewards.

The goal of this work is to identify the benefits and drawbacks of coordinating a formal AiR program as well as participating in in the program from the student-artist and faculty perspective. We recruited a variety of scientists and artists to collaborate in creating visual science communication products. We have defined sources for negative and positive experiences and outcomes from this AiR program using both qualitative and quantitative approaches. Further, we analyze responses from participants (student-artists and faculty) to specify areas for improvement and areas successful in producing a positive experience and outcome in an AiR program. We conclude this analysis with suggestions to enhance the benefits and outcomes of an AiR program and ways to minimize the difficulties faced by participants and program designers.

## METHODS

### Program development

#### Recruitment of student artists and faculty

Table 1 shows institutions of students and faculty that participated in the program over the last three years. We consider minority-serving status, NIH funding ranking, and geographical location (rural versus urban) for all student and faculty institutions. Since the program originated at Vanderbilt University/Vanderbilt University Medical Center, most students and faculty are located at this institution. If a student participated in multiple program years (N=2), they were counted twice and so on.

**Table 1.** Institutions of students and faculty. We consider minority-serving status, NIH funding ranking, and geographical location (rural versus urban) for all student and faculty institutions. Since the program originated at Vanderbilt University/Vanderbilt University Medical Center, most students and faculty are located at this institution. If a student participated in multiple years of the program, then they were counted twice (N=2).

#### Student-artists

In total, we have had 55 students participate in the AiR Program, which began in 2019. Students who participated in the program had a range of undergraduate experiences (Figure 1 A). However, most students were either sophomores or juniors. Students who participated in the AiR Program had a variety of majors and minors (Figure 1 B). If students were pursuing multiple majors or minors, each major and minor or a student was counted. The most popular majors were Biology or Biological Sciences, Medicine, Neuroscience, Health and Society, and Molecular and Cell Biology. The most common minors were art and chemistry.

**Figure 1.**
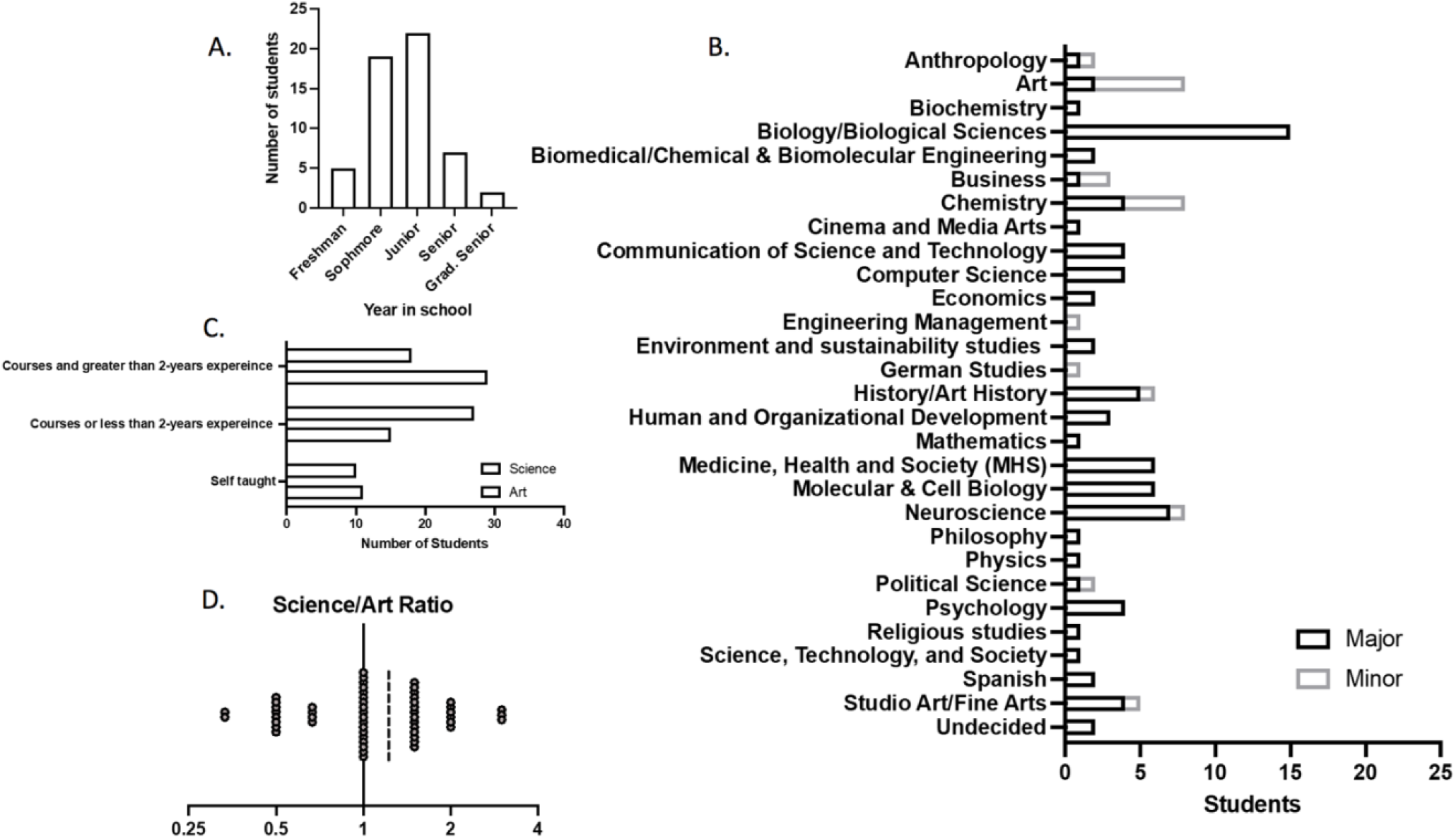
Demographics of AiR students. In total, we have had 55 students participate in the AiR Program, which began in 2019. A) Students who participated in the program had a range of undergraduate experiences. However, the majority of students were either sophomores or juniors. B) Students who participated in the AiR Program had a variety of majors and minors. If students were pursuing multiple majors or minors, each major and minor were counted equally. The most popular majors were Biology or Biological Sciences, Medicine, Neuroscience, Health and Society, and Molecular and Cell Biology. The most common minors were art and chemistry. C) Through a qualitative analysis from student applications, we explored and scored the experience in both art and science of the students who participated in the program. Students were given a score (1-3) for various experience levels in art and science. One indicated that the student was self-taught or had no experience. Two indicated that they had less than two years of practical experience (program, university-level research) experience or had only taken courses. Three indicated that they had taken courses with greater than two years of practical experience. Overall, many students took courses and had more than 2-years of university-level research experience in science and some courses or less than 2-years of program experience in art. D) To represent the balance of science versus art training, we examined the ratio of science over artist training. A value of 0.33 indicated much greater artistic experience over art experience, 1 indicating an equal amount of artistic and scientific training, and 3 indicating much greater science experience than the art experience. The mean value was 1.224 indicating greater scientific experience to artistic experience general for the AiR students.

#### Faculty

As we selected faculty, we were also interested in diversity, unique opportunities, and geographic location. We determined the faculty rank of the AiR faculty participants and found that most were full professors (Figure 2A). We also attempted to determine the faculty’s schools housed within various institutions (Figure 2B). The majority of faculty were in colleges or schools of medicine, primarily associated with medical institutions. Finally, based on appointment, we explored the faculty members’ department and field of study. The majority of faculty were either in Pathology, Microbiology and Immunology, or Medicine (Figure 3C).

**Figure 2.**
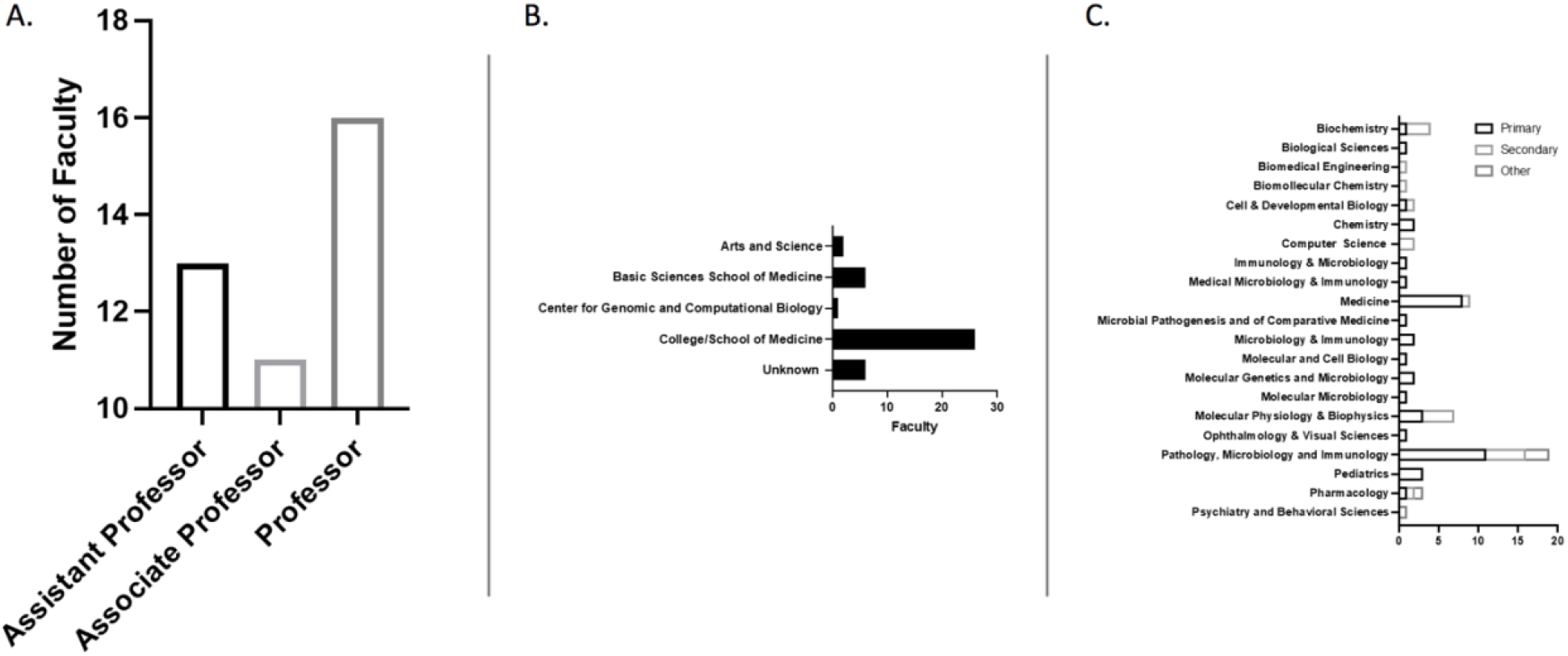
Demographics of AiR faculty. As we selected faculty, we were also interested in diversity, unique opportunities, and geographic location. A) We determined the faculty rank of the AiR faculty participants and found that most were full professors. B) We also attempted to determine the faculty’s schools housed within various institutions. The majority of faculty were located in colleges or schools of medicine, primarily associated with medical institutions. C) Finally, based on appointment, we explored the faculty members’ department and field of study. The majority of faculty were either in Pathology, Microbiology and Immunology, or Medicine.

**Figure 3.**
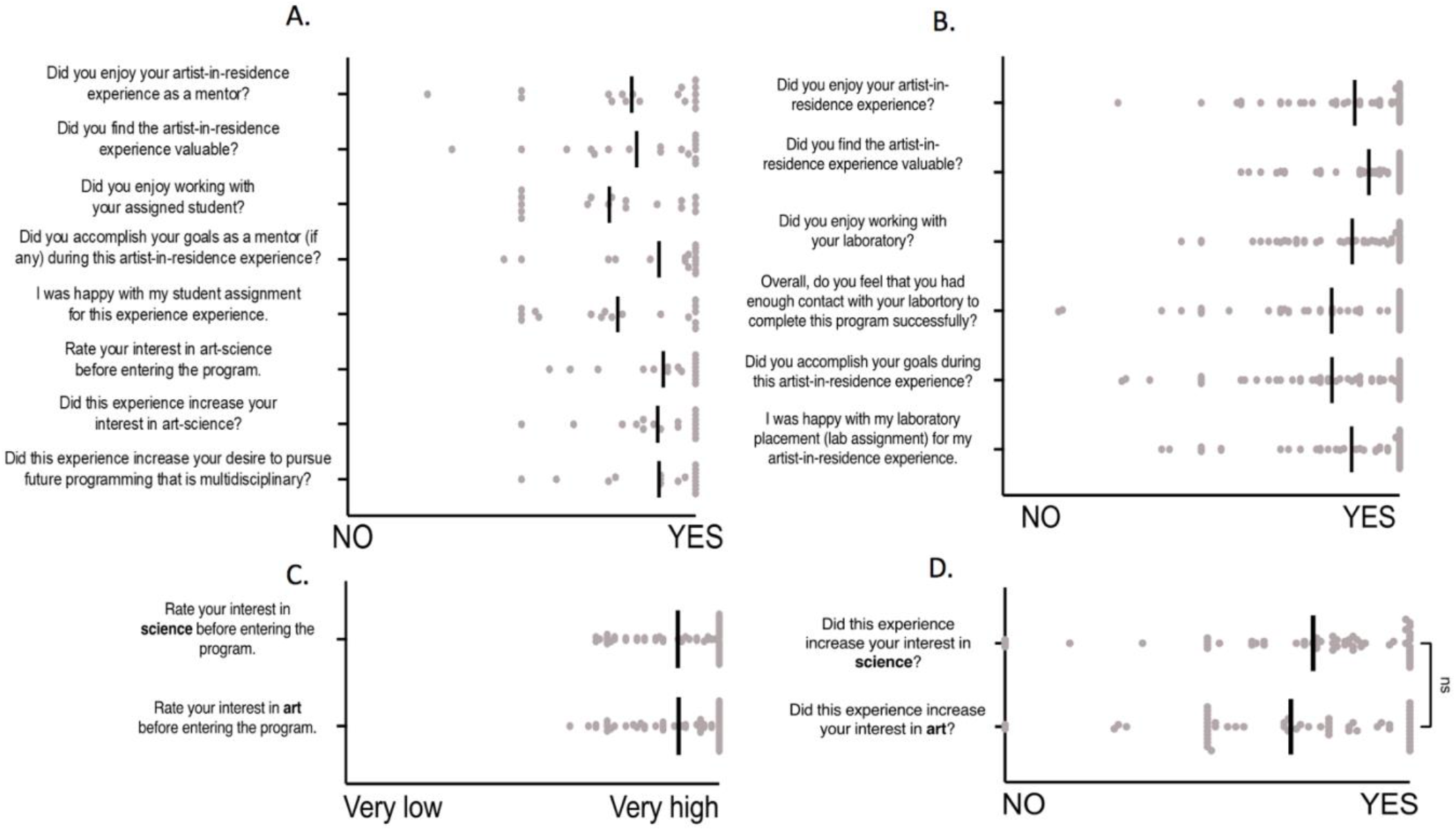
Review of experience in the program. Both students and faculty were asked a series of questions about their experience in the program. The answerers were collected on a sliding scale from 1-100 with (NO or Very Low equaling 0 and YES of Very High equaling 100). Each response is shown as an individual point, and the mean response is indicated with a black bar. Overall, both (A) students and (B) faculty responded “YES” to all questions posed. C) Students were also asked the rate their interest in science and art before entering the program. Students responded to being very interested in both art and science equally. D) When asked if participating in the program increased their experience, students overall answered “YES” to the experience, increasing their interest in science and art.

#### Study and survey design

This study was reviewed and approved by Vanderbilt University (Vanderbilt University IRB #220157). All participants had the opportunity not to complete the post-program survey. We have 100% of the students complete the post-program survey and 33.3% of faculty.

#### Quantitative analysis

For all data graphing, we used Prism 9.0 GraphPad software[32]. Three coders were used, and each coder was asked to examine the open-ended, de-identified survey response answers independently. To measure the agreement between coders, we used the NVivo coding comparison Query Criteria for each pair of coders that reports both a weighted and unweighted Kappa score. the Kappa scores were 0.42, 0.46, and 0.46 between the three reviewers indicating acceptable agreement of coders to continue with the analysis (more information available in supplemental methods).

#### Quantitively identifying student experience in art and science

Through a qualitative analysis from student applications, we explored and scored the experience in both art and science of the students who participated in the program (Figure 1 C). Application for the ArtLab program were shared through multiple networks. Likely the student who applied to the program had some interest in both art and science and therefore aren’t entirely random. The student applications were given a score (1-3) for various experience levels in art and science. One indicated that the student was self-taught or had no experience. Two indicated that they had less than two years of practical experience (program, university-level research) experience or had only taken courses. Three indicated that they had taken courses with greater than two years of practical experience.

#### Qualitative analysis

We used NVivo[33] for all qualitative data analysis. We set out to define (1) the AiR program experience from the perspective of student-artist versus faculty, and (2) outcomes of the program for student-artists versus faculty. We performed a constructive thematic analysis of student and faculty open-ended survey responses and coded open-ended responses into themes to answer these questions. The themes were informed by the program facilitators’ own experience and faculty responses. From this perspective and bias the theme generated about potential concerns related to “Interaction with program organizers,” “Responsiveness of program organizers when needed,” “Communication with the student,” and “Time to complete the project.” We combined the open-ended survey responses of students and faculty from a variety of prompts, including “Please explain your answer,” “What was your favorite aspect of the artist-in-residence experience?” “What was your least favorite aspect of the artist-in-residence experience?” and “What could make this experience more valuable in the future?” The author reviewed all open-ended responses to the post-program survey. Not all responses were required to be coded by the coders if they did not fit the themes. For more information, please see the supplementary methods.

## RESULTS

### Incoming experience levels of student artists

Overall, many students took courses and had more than 2-years of university-level research experience in science and some courses or less than 2-years of program experience in art. To represent the balance of science versus art training, we examined the ratio of science over artist training (Figure 1 D). A value of 0.33 indicated much greater artistic experience over science experience, 1 indicating an equal amount of artistic and scientific training, and 3 indicating much greater science experience than art experience. The mean value was 1.224 indicating greater scientific experience to artistic experience general for the AiR students.

### Experience in the program

Both students and faculty were asked a series of questions about their experience in the program. The answers were collected on a sliding scale from 1-100 with (0 meaning “NO” or “Very Low” equaling and 100 meaning “YES” or “Very High” depending on the question). Each response is shown as an individual point, and the mean response is indicated with a black bar. Overall, both students and faculty overwhelming responded “YES” to all questions posed (Figure 3A and B). Students were also asked the rate their interest in science and art before entering the program (Figure 3C). Students responded to being very interested in both art and science equally. When asked if participating in the program increased their experience, students overall answered “YES” to the program increasing their interest in science and art (Figure 3D).

Within the survey, we also examined Likert scale data from the final program survey that explored student and faculty satisfaction with various aspects of the program (Figure 4). This number of responses for each prompt is reported via a heat map. Overall, students reported being “very satisfied” with all provided prompts listed in Figure 4. Faculty responses were more varied, particularly regarding “The remote (virtual) experience,” “Interaction with program organizers,” “Responsiveness of program organizers when needed,” “Communication with the student,” and “Time to complete the project.” We used these prompts to inform a constructive thematic qualitative analysis conducted on the open-ended survey questions from both students and faculty.

**Figure 4.**
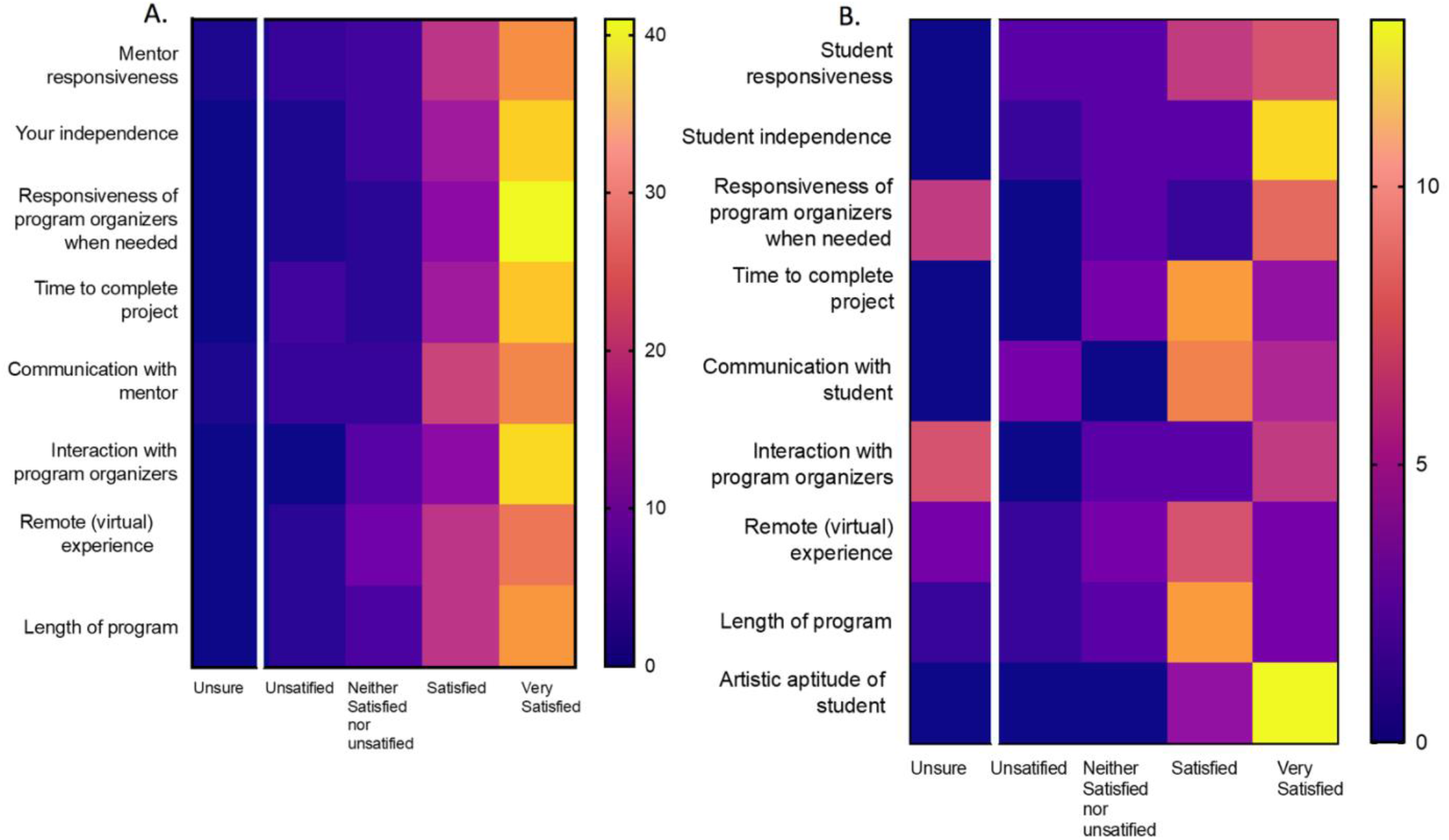
Satisfaction data. We examined Likert scale data from the final program survey that explored (A) student (N=55) and (B) faculty (N=17) satisfaction with various aspects of the program. This number of responses for each prompt is reported via a heat map, with purple indicating fewer responses and yellow indicating a more significant number of responses. Overall, students reported being “very satisfied” with all provided prompts. Faculty responses were more varied, particularly regarding “The remote (virtual) experience,” “Interaction with program organizers and Responsiveness of program organizers when needed,” “Communication with the student,” and “Time to complete the project.” We used these prompts for a constructive thematic qualitative analysis conducted on the open-ended survey questions from both students and faculty presented in Figure 6.

### Word clouds as an overview of open-ended responses

This led us to explore a deeper understanding of the student versus faculty experience. We, therefore, collected all open-ended responses from both students and faculty for analysis. Before beginning a directed content analysis, we generated a word cloud of open-ended survey responses from students and faculty (Figure 5). Word clouds are used in various contexts as a means to provide an overview by distilling text down to those words that appear with highest frequency. The top 100 words with a minimum length of 3 letters were plots in a word cloud. All responses from students (A) and faculty (B) were coded and grouped based on either being from a student or a faculty participant. In viewing the word clouds, an overview of the words that occur most often within the text is visualized. These words included

**Figure 5.**
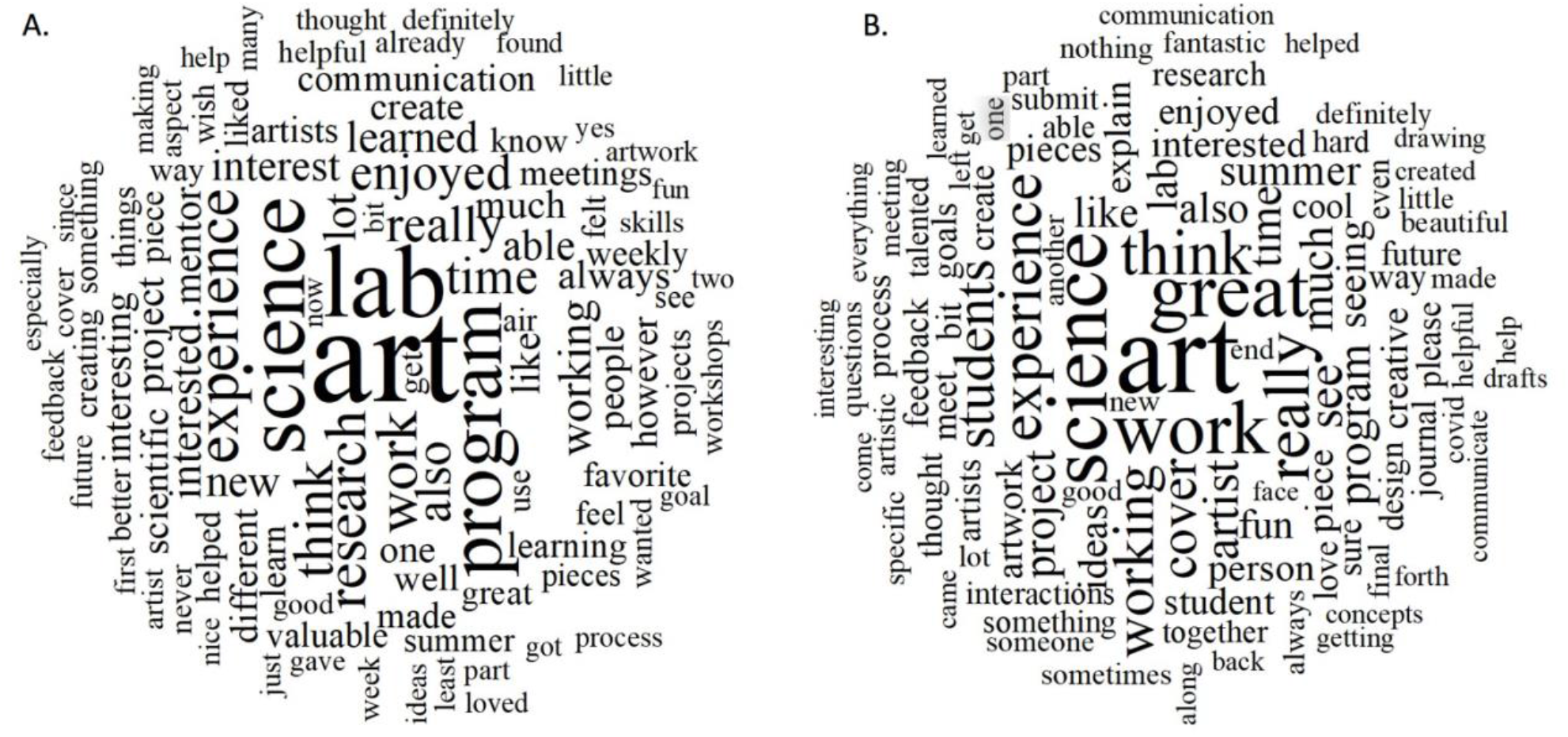
Word cloud of open-ended responses. Word cloud of open-ended survey responses from (A) students and (B) faculty. The top 100 words with a minimum length of 3 letters were plots in a word cloud. All responses from students and faculty were coded and grouped based on either being from a student or a faculty participant. The purpose of this coding was to explore the experience of both students and faculty. Before conducting a constructive thematic analysis, this up-biased representation provides an overview of the responses.

### Informed thematic analysis for lower-performing satisfaction areas among faculty

Based on the satisfaction analysis from Figure 4, we wanted to better understand the areas that did not exceed expectations. The four themes investigated as part of the program experience included communication between program participants, program design and implementation, time or length, and the virtual or remote nature of the program. These themes were informed based on results from Figure 4. We compared the number of responses as well as the amount of coverage each theme had among all responses, or percent coverage, to minimize the disparity between the number of responses between students and faculty. All percentages below refer to the percentage of all open-ended text responses from faculty or students that represents that theme. Low percentages indicate that there are fewer comments about that theme, while higher percentages indicate that there was more comment about that theme.”

#### Communication between program participants

The first area we explored with the thematic analysis was communication between program participants (Figure 6 A). We flagged any comment related to program communication between students, faculty, or student-faculty partners. Coders were instructed not to include comments on science communication for a public audience to better understand the communication between the student and faculty partnership. Overall, there were 318 coding references for students (20.96%, coverage of all open response text from students) and 85 coding references for faculty (28.93%, coverage of all open response text from faculty) related to communication (Supplemental Figure 4 and 5). There was a balance of both positive and negative codes from both student and faculty about communication between participants but no significant difference (Supplemental Figure 1).

**Figure 6.**
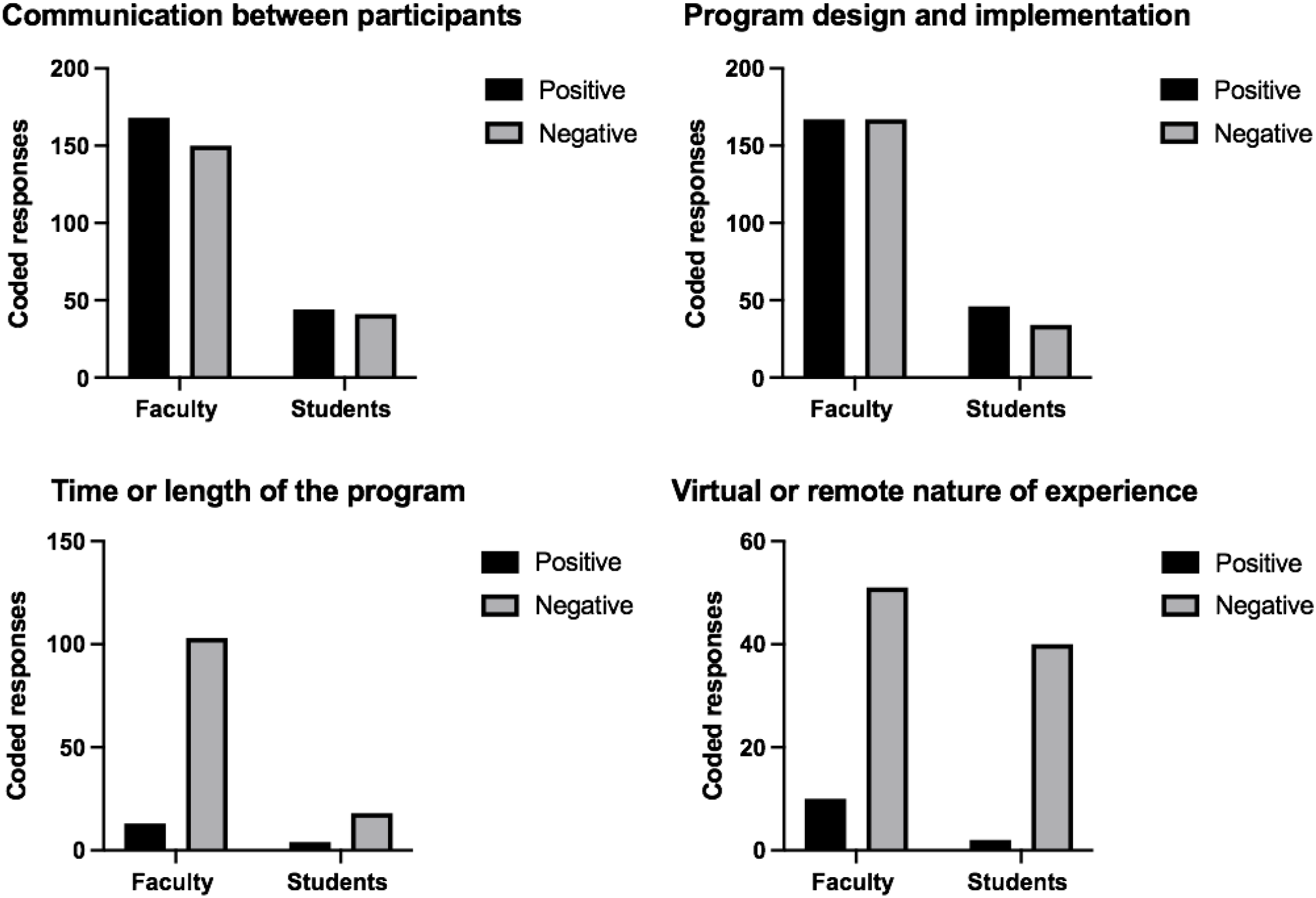
Informed thematic analysis on program experience between students and faculty. We analyzed all codes within each theme for differences in the positive or negative responses for that theme. For each theme, a chi-squared analysis was performed using the raw code values within each group. For A) “communication between participants,” B) “program design and implementation,” C) “time of the length of the program,” and D) “virtual or remote nature of experience,” there is no difference between students and faculty in the percentage of responses that were coded at being relevant for the program experience. Student and faculty experiences trend similarly; There was a balance of positive and negative codes for A) “communication between participants,” and B) “program design and implementation,” and a similarly negative perspective on the “time or length of the program” and the “virtual or remote nature of the experience.”

For student responses, 168 coding references were positive (10.83%), and 150 were negative (10.44%). Examples of positive student codes include, “My discussions with my mentor have been very efficient,” “I feel that my contact person has been perfect with communicating,” and “I thoroughly enjoyed being paired with a lab and our weekly meetings to see what the other artists were up to.” Some examples of negative student comments include, “I wish I had more communication with my mentor,” “Besides the lab I was working with, I didn’t have any people to ask questions because I had never done a residency before, so it was feeling around in the dark to find something that worked,” and “Discussing and deciding on the subject of the piece pushed the creation of the piece back significantly, and I believe the execution of my final piece suffered because of it.”

For faculty, 44 coding references were positive (17.04%), and 41 were negative (12.94%). Examples of positive faculty responses include, “We got along great, and I think it came through in the output,” “She did a phenomenal job of integrating scientific paper abstracts into art ideas and went above and beyond in soliciting feedback and generating drafts,” and “She provided updates regularly and responded well to suggestions. She also was eager to learn about this project and asked questions when needed.” Some examples of negative faculty responses related to communication include, “Aside from our initial interaction by phone, there was not much follow-up or dialogue,” “It was sometimes hard to schedule meetings with the artists,” and “Trying to schedule with the artists, it seems like they had many other demands on their time.”

#### Program design and implementation

The second area that we examined relating to the program experience was the program’s design and implementation (Figure 6 B). This included comments that relate to program organizers, program organization, including learning goals, objectives, emails, approach, guidance, and other organizational elements. Overall, there were 334 coding references for students (24.47%) and 80 coding references for faculty (23.44%) related to the program design and implementation. There was a balance of positive and negative codes from both students and faculty about program design and implementation but no significant difference.

For students, 167 coded references were positive (13.74%), and 167 coded references were negative (10.95%). Examples of positive comments include, “Filling the survey provided by ___ also helped me clear my thoughts and inspired me a lot,” “having the opportunity to have my artwork printed and included in an exhibition for the year is incredibly rewarding and something that I hope to continue in the future,” “I thoroughly enjoyed being paired with a lab and our weekly meetings to see what the other artists wer e up to.” Some examples of negative comments related to program design and implementation include, “I just wish the beginning of it was more coordinated. Besides the lab I was working with, I didn’t have any people to ask questions because I had never done a residency before, so it was feeling around in the dark to find something that worked,” “However, I wish there was a clear publication or project selected from the beginning of the project. Discussing and deciding on the subject of the piece pushed the creation of the piece back significantly, and I believe the execution of my final piece suffered because of it,” and “I was also expecting for there to be more technical workshops learning techniques in software.”

For faculty, 46 coded references were positive (12.68%), and 34 coded references were negative (10.76%). Some examples of positive comments from faculty include, “the best part was seeing our work from an artistic viewpoint and developing an abstract drawing that would present our work in a creative and aesthetic way,” “I made sure to engage multiple lab members in the process to make the experience as rich as possible for everybody,” and “I most enjoyed the brainstorming and concept sessions.” Some examples of negative comments related to program design and implementation include, “having a little bit of structure to go back and forth on ideas and drafts could have been helpful,” “artist not seeming to have full guidance and direction,” and “having a better understanding of the expectations might have helped me …, for example, if I knew that it was expected to go back and forth 4-6 times on drafts then I would know that giving feedback five times is reasonable and not too much to ask.”

#### Time or length of the program

Next, we exampled the theme of time or length of the program (Figure 6 C). This theme related to all comments that mention the scheduling, timing, not finishing a project due to time limitations, time management, etc. Overall, 116 coded references for students (7.39%) and 22 coding references for faculty (4.56%) related to the program time and/or length. There were more negative codes from both students and faculty about communication between participants but no significant difference between students and faculty in terms of positive or negative codes.

For students, 13 coded references were positive (0.77%), and 103 coded (6.65%) references were negative. Examples of positive comments include, “I enjoyed the way of building up a set of works from continuous efforts in a given amount of time.” Some examples of negative comments include, “Overall I enjoyed the program, but feel that it was too short to do my best work,” “This would have been an incredibly rewarding experience; my only regret is taking on too much this summer, so I did not have the time to devote to this program,” and “… the stress of completing the weekly surveys and watching lectures made everything seem rushed.”

For faculty, four coded references were positive (0.68%), and 18 coded references were negative (3.95%). Some examples of positive comments include, “It was great practice in scientific communication for me, and didn’t take up too much time,” and “it was great to do this over the summer.” Some examples of negative comments related to the length of the program include, “[my least favorite part] How short it was,” and “Time flew by quickly summer, and I wish we could have had a bit more time to work together.”

#### Virtual or remote nature of experience

Finally, the last theme that we examined as a part of the program experience was the virtual or remote nature of the program (Figure 6 D). Comments coded to this theme included any mention of being in-person, working remotely, or interacting virtually. Overall, there were 61 coded references for students (2.76%) and 42 coded references for faculty (8.53%) related to the program’s virtual or remote nature. There were more negative codes associated with the virtual or remote nature of the experience from both student and faculty about communication between participants but no significant difference. However, it was trending towards more negative faculty responses than students.

For students, ten coding references were positive (0.64%), and 51 coded (2.71%) references were negative. An example of a positive student response was, “I thought the program was well adapted to a remote-only environment.” An example of a negative comment about the virtual experience included, “The only thing that influences my experience is the technical inconvenience of communication sometimes due to internet control in China,” “However, being remote has made it difficult to find inspirations for the project,” and “However, I think that an in-person experience in the future would be even better!”

For faculty, two coding references were positive (0.38%), and 40 coding references were negative (8.33%). An example of positive comment about the virtual experience was, “But overall, it was a fantastic experience, and converting to remote still worked great!” Some examples of negative comments related to the virtual experience include, “I think it was a bit hard to do this remotely, but I can understand why it was necessary,” “The fact that most of it was virtual [was my least favorite part],” and “[my least favorite part was] not having her nearby or getting to meet her in person before the program started.”

### Thematic analysis of benefits for artist-scientist partnership

In addition to running a thematic analysis on the program experience, we performed a parallel investigation to explore program outcomes. The four themes related to program outcomes that were coded include mentoring experience, way of thinking, learning something new, and science communication to the public or general audience. Again, we compared the percent coverage of each theme to minimize the disparity between the number of responses between students and faculty.

#### Mentoring experience

The first theme coded for part of the program outcomes was the mentoring experience (Figure 7A). This included comments related to either a student or a faculty mentoring situation in a programmatic sense. This did not include specific statements about mentoring styles but focused more on mentorship. Overall, there were 339 coding references for students (22.5%) and 151 coding references for faculty (42.49%) related to the mentoring experience. There were more positive codes from both students and faculty about the mentoring experience. There was no significant difference between students and faculty regarding positive or negative codes.

**Figure 7.**
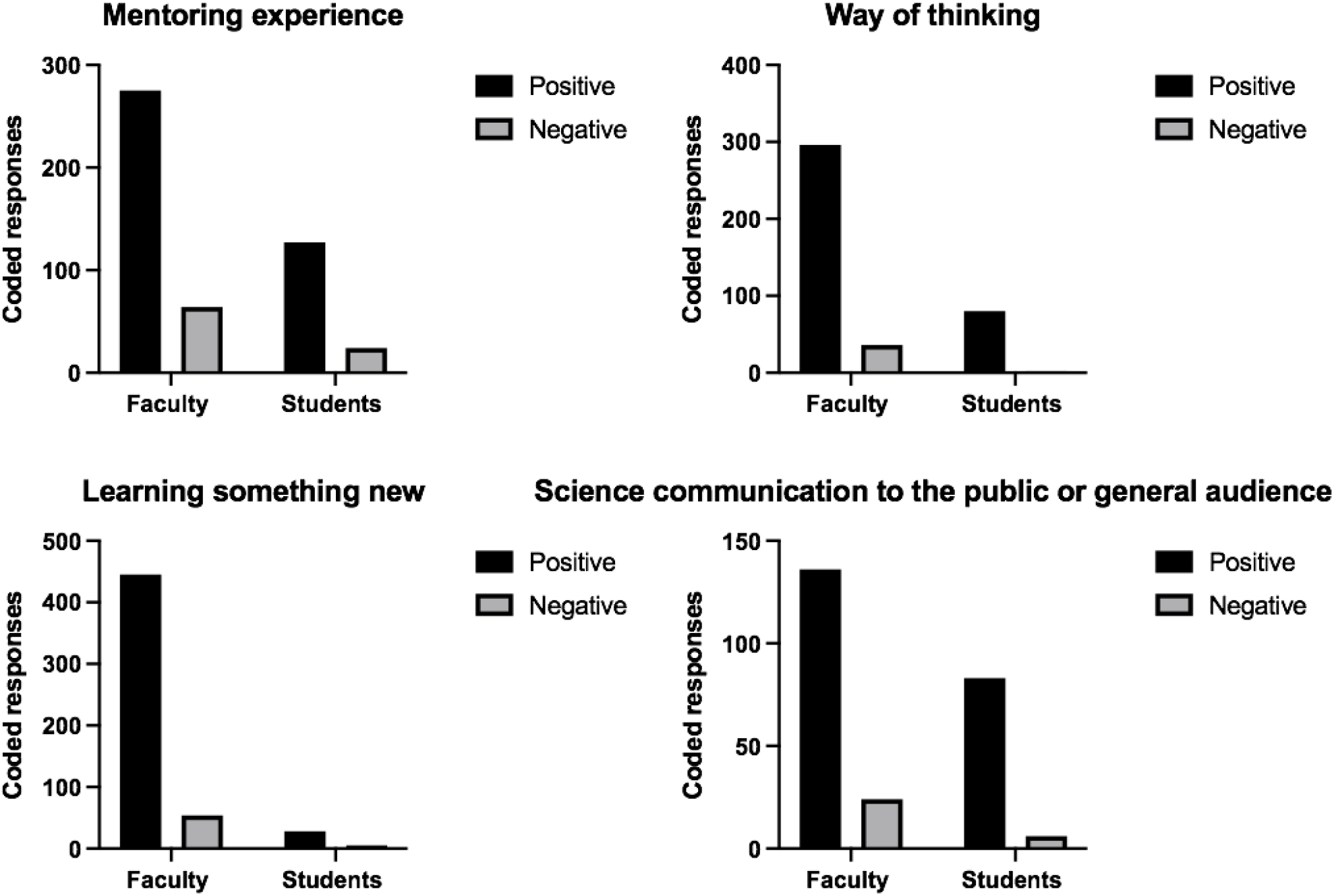
Thematic analysis of program outcomes for students and faculty. We analyzed all codes within each theme for differences in the positive or negative responses for that theme. For each theme, a chi-squared analysis was performed using the raw code values within each group. For A) “mentoring experience,” C) “learning something new,” no significant difference was observed between student and faculty perspectives. There were far more positive outcomes related to “mentoring experiences” and “learning something new” for students and faculty. For B) “way of thinking,” D) “science communication,” there was a significant difference (B) or a trend towards a significant difference between students and faculty.

For students, 275 coded references were positive (15.51%), and 64 coded references were negative (7.1%). Some examples of positive comments are, “My lab was incredibly enthusiastic and welcoming,” “I enjoyed working in conjunction with the ___ lab to explore where my two interests merge,” and “I enjoyed getting to know ___ and ___.” Some examples of negative comments related to mentoring included, “The only thing keeping me from loving this experience 100% was the absence of my lab PI throughout my time in the program,” and “I felt that it was sometimes unclear how the PI wanted to use my work and what the goals were for a final product.”

For faculty, 127 coded references were positive (32.39%), and 24 coded references were negative (10.5%). Some examples of positive comments include, “It was a blast working on art projects with such a creative and kind person,” “___ came in with so many ideas and asked really insightful questions about the science to represent everything correctly,” and “___ and I built a really positive rapport that went beyond our specific project. I felt like it was a really great mentoring experience, and we plan to keep in touch!” Some examples of negative comments related to mentoring included, “Aside from our initial interaction by phone, there was not much follow-up or dialogue,” and “I only partially mentored the student. I was more of a liaison between her and the graduate students and postdocs.”

#### Way of thinking

The second theme coded for was “way of thinking,” which was used to indicate comments related to how the experience led to a change in the student or faculty members’ way of thinking related to science, art, or in general (Figure 7 B). Overall, there were 332 coded references for students (23.48%) and 82 coded references for faculty (25.05%) related to a change in thinking. There were more positive codes from both students and faculty about the theme and there was a significant difference between student and faculty. Overall, faculty had a more significant percentage of positive codes for the theme and fewer negative codes than students.

For students, 296 coding references were positive (20.56%), and 36 coding references were negative (3.03%). Some examples of positive comments related to the way of thinking included, “I enjoyed exploring the art side of science,” “I really think that I was encouraged to explore and try new things in the program, and I’ve learned many new art techniques,” and “it was challenging but fun to figure out how to represent them visually.” An example of a negative comment relating to the “way of thinking” was, “Thinking up creative ways to communicate research accurately is a difficult task, and I would have to sacrifice either creativity or accuracy or ease of understanding.”

For faculty, 80 coding references were positive (24.44%), and two coding references were negative (0.61%). Some examples of positive comments related to ‘way of thinking” are, “It was great to think creatively with a true artist about visual scientific communication,” “these are very talented students that introduced new mediums and ideas of how to communicate science to others outside of my circle,” and “It was fun to explain the project that I’ve been working on to someone with no science background, and then to see it brought to life by the artist.” An example of a negative comment related to “way of thinking” was, “I already value and search for [the intersection of art and science] in our research efforts.”

#### Learning something new

The next theme that we explored as a part of program outcomes was “learning something new” (Figure 7 C). We used this code to tag any response where the individual indicated that they had learned or experienced something new or unique related to art or science from the program. Overall, there were 499 coded references for students (27.58%) and 33 for faculty (10.73%) related to learning something new. There were more positive codes from both students and faculty about “learning something new,” and there was no significant difference between student and faculty.

For students, 445 coded references were positive (23.23%), and 54 coded references were negative (4.53%). Some examples of positive comments related to learning something new included, “It was interesting because I learned more about the ___ lab’s work by reading the manuscripts while looking up terms on my own,” “I really think that I was encouraged to explore and try new things in the program, and I’ve learned many new art techniques,” and “Through the experience in AiR program, [I] not only got to know biological knowledge and research works but also learned so many new art techniques.” Some examples of negative comments related to learning something new included, “I wish I had learned more overall,” and “The workshop meetings sometimes felt very separate from what we were doing in the program, and I sometimes struggled to make a connection to my own work.”

For faculty, 28 coded references were positive (9.82%), and five coded references were negative (2.14%). Some examples of positive comments from faculty about learning something new included, “it was interesting to hear an artistic perspective on science,” “[My favorite part was] finding a way to relate to someone who is not in my field,” and “Having that creative vision and artistic skill to produce cover art is something I’d never be able to do on my own.” An example of a negative comment related to learning something new was, “My interest [in art-science] was already high, it didn’t increase.” Comments similar to these often corresponded to those with some pre-existing art-science background.

#### Science communication to the public or general audience

Finally, we explored the theme of “science communication to the public or general audience” (Figure 7 D). We used this code to tag any comment or response that relates to a positive impact that the program made on the student or faculty members’ ability to communicate or understand science. Overall, there were 160 coded references for students (13.71% coverage) and 89 coding references for faculty (26.22% coverage) related to science communication. There were more positive codes from both students and faculty about the “science communication to the public or general audience,” and there was a trend towards the difference between student and faculty. Overall, faculty had a greater percentage of positive codes for “science communication to the public or general audience” and fewer negative codes than students.

For students, 136 coding references were positive (11.85%), and 24 coding references were negative (1.9%). Some examples of comments related to positive program outcomes around the theme of science communication included, “I loved exploring scientific communication and illustration” and “I liked working with a lab and figuring out how to communicate the scientific concepts in a creative way.” An example of a comment negatively related to science communication was, “I thought we would learn more about how to effectively communicate science through art, which we may have briefly touched on in some sessions but not completely.”

For faculty, 83 coding references were positive (24.1%), and six coding references were negative (2.11%). Some examples of positive comments from faculty that were related to science communication included, “It was great to think creatively with a true artist about visual scientific communication,” “Working with ___ to design cover art was a fun and incredibly valuable experience,” and “This was a great experience that resulted in a beautiful piece of art that encompassed the major concepts of our research.” An example of negative comments related to science communication was, “It was fun, and I am excited to have this art piece, but whether we’ll end up using it professionally (journal cover, etc.) I’m not so sure.”

## DISCUSSION

The goal of this paper is to review three years of post-programmatic data from a virtual AiR program to (1) determine what the benefits are for both faculty and student artists, (2) improve the design and implementation of the program, and (3) develop best practices for art-science based partnerships. Our programmatic evaluation is particularly relevant for undergraduate student artists and faculty in the biomedical sciences. Our findings suggest an opportunity for alterations within the program to enhance the program experience for both students and faculty. Furthermore, we have found that the program experience was similar between students and faculty. Overall, program outcomes were positive by all participants, which suggests the benefits of this AiR program, including experiencing mentorship, altering one’s way of thinking, learning new skills, and implementing science communication. However, our results suggest that particularly for “way of thinking” and “implementing science communication,” faculty may benefit more.

As we began to explore our data, we became interested in defining the program experience from student-artist versus faculty. In many cases, both students and faculty have similar positive and negative experiences during the artist in residence program. For instance, there was a significantly higher number of negative responses regarding the time or length of the program and the virtual and remote nature of the program than positive responses from both students and faculty. Because not all of our theme demonstrated higher negative responses, we do not believe that the larger number of negative responses for themes like “time or length of the program” and “virtual or remote nature of the experience” is simply a survey error. These findings suggest the need to modify the program’s virtual nature, whether that be shifting to a hybrid or fully-in person experience. If the program were to remain virtual, then additional effort should be put on demonstrating the benefits of this modality, particularly for faculty. Altering the length of the program should also be addressed to better the overall experience for both students and faculty.

Similarly, as we reviewed the open responses, we were interested in better defining what the students and faculty valued program outcomes. We created a thematic analysis based on the open-ended responses related to mentoring, way of thinking, learning new skills, and science communication. In many instances, the faculty and students have similar values of the program outcomes. For example, a significantly higher percentage of positive responses regarding program outcomes of “mentorship experience’ and “learning something new” than negative responses from both students and faculty. However, some examples of student and faculty perspectives on program outcomes differed. For instance, student participants had more positive responses about learning something new from the program than faculty members. However, the most surprising aspect of our analysis suggests that for both “way of thinking” and “science communication to the public or general audience,” there may be more significant beneficial gains for faculty compared to students. This information could be useful in future designs of art-science programs by customizing the programming for these distinct benefits for each participant group.

From this analysis future program development and targeted outcomes can be improved. This includes specific modifications to the program experience, including transforming the program into a hybrid nature, with options and support for in-person components. This modification could give more insight into an optimal way of collaborating versus restricting to virtual or in-person since both natures have positive and negative responses. Based on responses, lengthening the program or putting less demand on deadline for the final product could increase positive experience reporting. However, there could also be concerns about not specifying program outcomes goals and creating deadlines. Responses regarding communication suggest subjectivity based on the student-faculty matching and could be improved with program design, implementing more communication methods, and increased collaboration with program directors to ensure efficient communication. Responses for program design also request more clarification of the overall process of the program and final product expectations, which could increase positive experience reporting.

There are several limitations to this study that readers must take into consideration. First, as previously mentioned, the themes developed for the constructive thematic analysis were based on the program directors’ first-hand experience. Therefore, the themes might introduce confirmation bias based on the program director’s experience and subjective perspective. Because we used these defining themes to guide the data analysis, there may be more or parallel information within the data set that was not collected or represented in the final analysis. We attempted to mitigate this bias by using three coders. Another major limitation of this study was selection bias because the students and faculty that participated in this program have a pre-established interest in merging art and science. This is particularly evident in Figure 3, where students were asked the rate their interest in science and art before and after entering the program. Because the student interest was extremely high before entering the program, we reached a ceiling effect where there was less than expected growth in interest after participation. Finally, It is unclear why faculty participation was only 33% for the post-program survey. However, we feel that these responses represent the overall pool of faculty participants. Future studies conducted by those outside of the program and involving student-faculty pairs that do not have pre-established interest in art and science intersection may expand the impact of this study.

Program outcomes of “mentorship experience,” “way of thinking,” “learning something new,” and “science communication” were beneficial among both students and faculty. Still, our analysis suggests that for “way of thinking” and “science communication,” there could be even more significant gains for faculty as compared to students. There is a need to improve the skills and provide training opportunities for faculty to talk about their research in effective and compelling ways. Many efforts are underway to provide faculty with experience speaking about their research to the public. The AiR experience might be one avenue for science communication training opportunities that also provides a tangible visual science communication product for the faculty member.

Overall, this analysis summarizes the AiR experience from the perspective of both students and faculty to understand how to design the program and the benefits for both students and faculty. From this analysis, we can take clear steps to maximize the program experience, and they would also be likely to enhance the program benefits further. For instance, negative comments regarding the mentorship experience included communication issues that we could address by adjusting the program design on the communication suggested above. We have defined promising benefits of running an AiR program, diagnoses of difficulties in an AiR program, insight on bettering program design and implementation, and furthering future developments beneficial outcomes of practices of art-science partnerships.

## AVAILABILITY OF DATA AND MATERIALS

A PDF of the overview data and analysis is available upon request. Survey data may be made available upon reasonable request.

## ACKNOWLEDGMENTS

We thank the faculty and 55 students that participated in our programming.

## ETHICS DECLARATION, PROJECT TITLE

## ETHICS APPROVAL AND CONSENT TO PARTICIPATE

Yes

## CONSENT FOR PUBLICATION

Yes

## FIGURES

*(Figure legends included in manuscript file)*

