## Supplemental for "Differences in self-reported benefits for student-artist versus faculty experiences in a virtual artist-in-residence program"

### SUPPLEMENTARY INFORMATION

#### Supplemental Methods

We developed two sets of codes. We labeled comments as either positive or negative for both coding approaches.

- ***Program design.*** The first set of codes was based on the satisfaction data and the responses of faculty that indicated that they seemed not to meet the highest satisfaction. The codes for the first set of themes are:
  - **Communication.** We used this to code anything that mentions or discusses communication between the student and the mentor, impacting the program experience.
  - **Virtual Experience.** This code was used when a response mentions issues related to the experience between virtual or remote or predicting an in-person experience.
  - **Program organization and implementation.** We expanded this code to include responsiveness and communication with program organizers and all aspects of program organization, including goals, objectives, emails, approach, etc.
  - **Time.** This code was used when a response talks about the time required to complete the project, either relating to having enough time or not enough time to complete the project as intended.
- ***Program outcomes.*** The second code set specifically exampled the program's benefits for both faculty and students. The codes of this set include:
  - **Way of thinking.** We used this code to tag any response related to how the experience led to a positive change in the student or faculty members' way of thinking, either related to science, art, or general.
  - **Mentoring.** We used this code to tag any response related to how the experience led to a positive or negative mentoring opportunity either as a student or a faculty member.
  - **Communicating.** We used this code to tag any response related to the program's impact on student or faculty members' ability to communicate or understand science.
  - **New Things.** We used this code to tag any response where the individual indicated that they had learned or experienced something new or unique related to either art or science from the program.

**Coding of open-ended responses.** Three coders were used, and each coder was asked to examine the open-ended, de-identified survey response answers independently. The principal investigator provided coders with a coding guide. Coders were asked to read through the open-ended responses and code both sets of thematic categories corresponding to that response, if appropriate. A comment could be coded for no theme or multiple themes. Finally, the coders were asked to indicate whether the response was positive or negative. The results were compiled and reviewed by the principal investigator directing the project.

To measure the agreement between coders, we used the NVivo coding comparison Query Criteria for each pair of coders that reports both a weighted and unweighted Kappa score. A Kappa measurement accounts for the chance-adjusted measure of agreement for any number of cases, categories, or raters [1–3]. Kappa values can range from -1.0 to 1.0. The lower end of the scale, -1.0, indicates perfect disagreement below chance. 0.0 indicates agreement equal to chance. The higher end of the scale, 1.0, indicates excellent agreement above chance. Typically, kappa values less than .40 are "poor," values from .40 to .75 are "intermediate to good," and values above .95 are "excellent." For all coding, the Kappa scores were 0.42, 0.46, and 0.46 indicating acceptable agreement of coders to continue with the analysis. The supplemental table shows the number of responses that were coded as being related to each theme and if they were positive or negative for students and faculty. The relative frequency (percentage of all codes) of comments and if they were positive or negative for each theme is provided. We migrated data from NVivo[4] to GraphPad Prism[5] for graphical representations.

**Supplementary Document 1.** Coding guide.
Available for download: https://vanderbilt.box.com/s/uk1rgqbbqg2akzq0jq8ma0ug4e3vy7jv

### Supplemental Tables and Figures

**Supplementary Figure 1. Comparing student to faculty responses.** To determine if students and faculty responded differently to the prompts about their experience in the program, we graphed student and faculty responses identical to Figure 3 on the same graph. There was no significant difference between faculty and staff responses to survey questions.

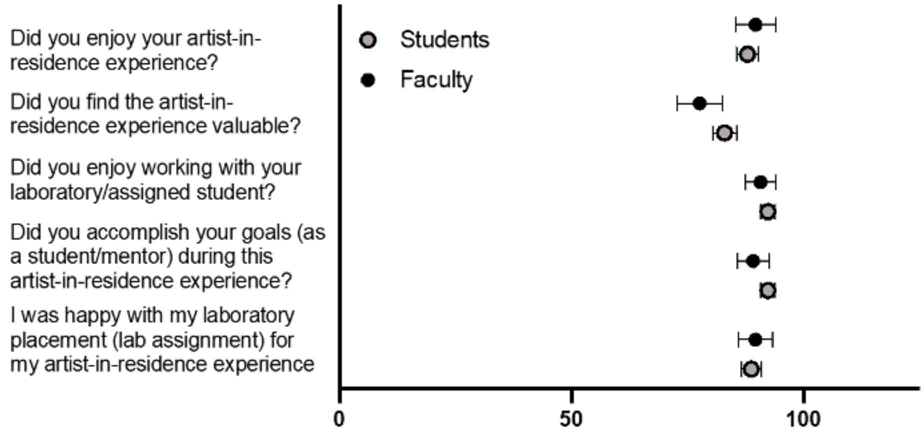


**Supplementary Figure 2.** Coding themes.

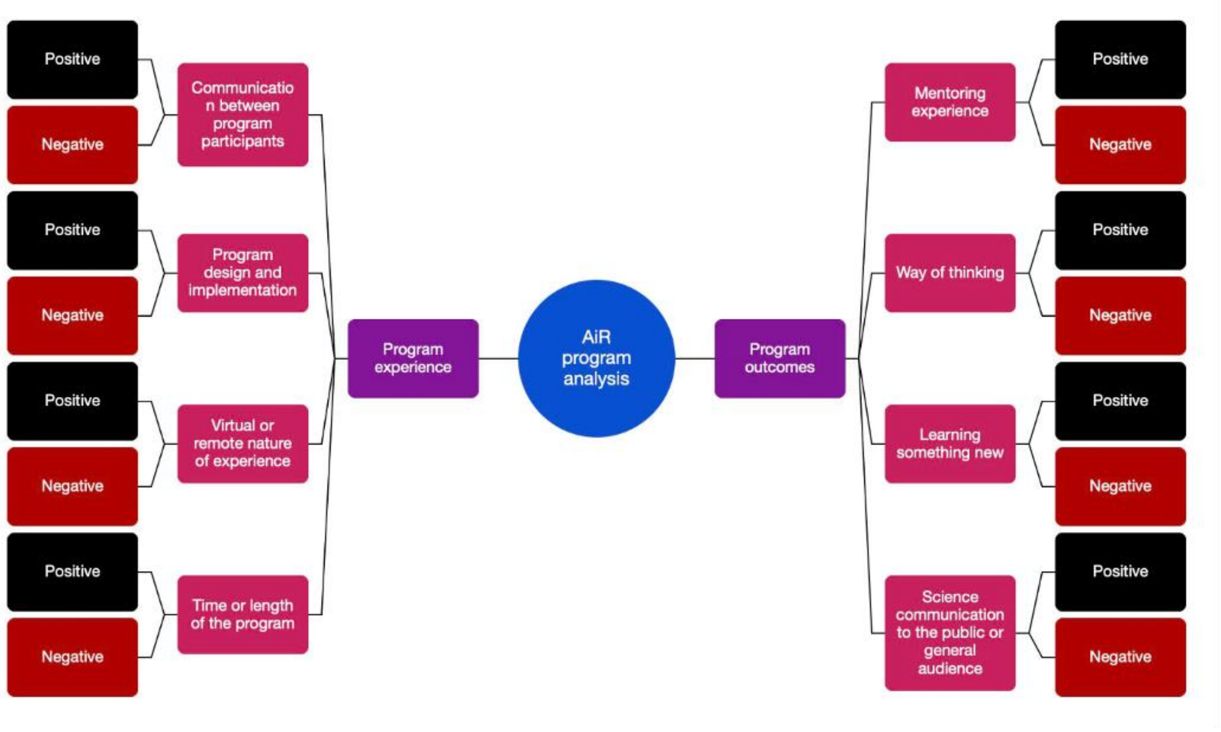

**Supplementary Figure 3.** Inter-coder reliability Kappa scores.

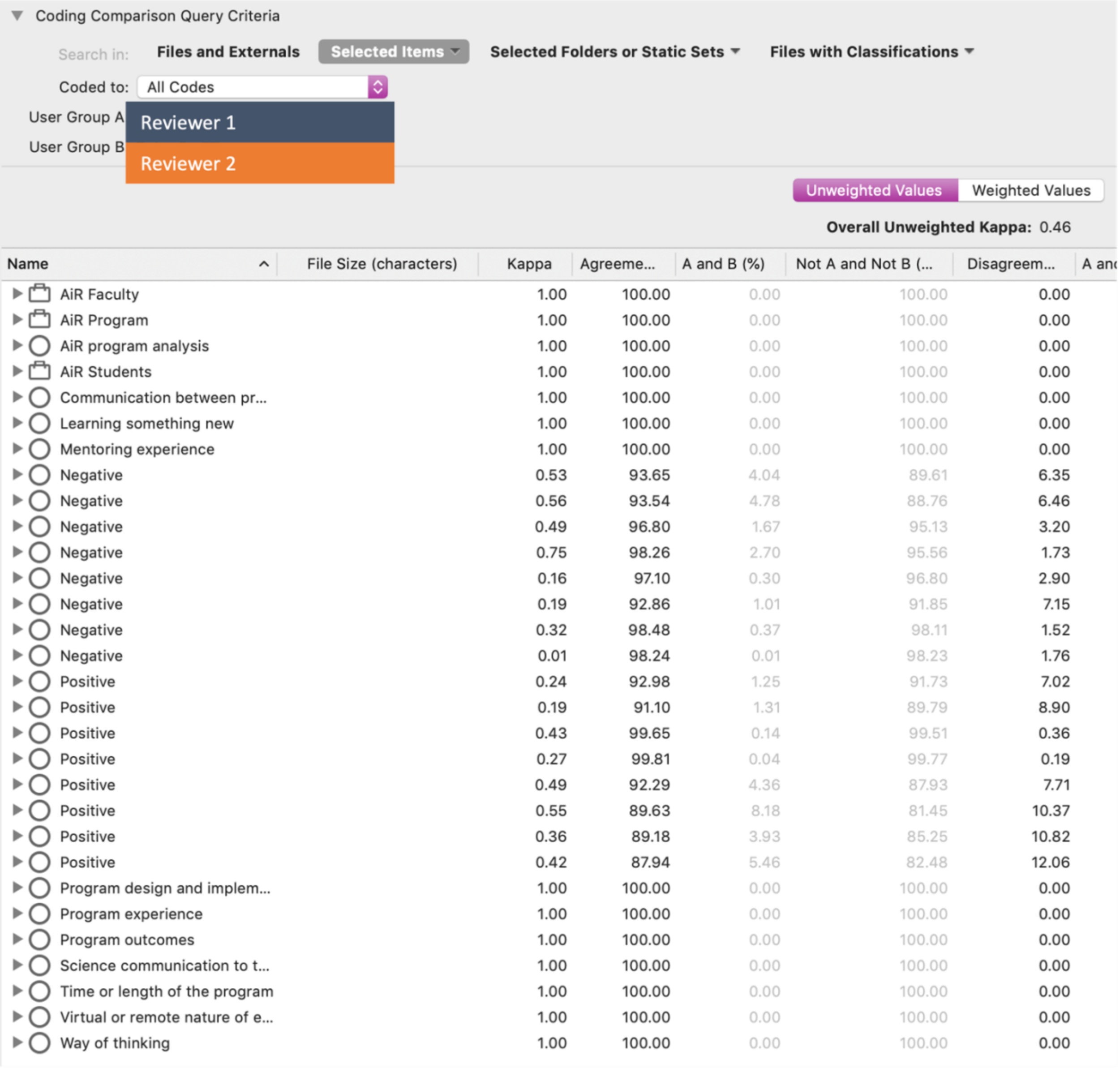

Coding Comparison Query Criteria

Search in: Files and Externals | Selected Items | Selected Folders or Static Sets | Files with Classifications

Coded to: All Codes

User Group A: Reviewer 1

User Group B: Reviewer 2

Unweighted Values | Weighted Values

**Overall Unweighted Kappa:** 0.46

| Name | File Size (characters) | Kappa | Agreeme... | A and B (%) | Not A and Not B (... | Disagreem... | A and |
|---|---|---|---|---|---|---|---|
| AiR Faculty | | 1.00 | 100.00 | 0.00 | 100.00 | 0.00 | |
| AiR Program | | 1.00 | 100.00 | 0.00 | 100.00 | 0.00 | |
| AiR program analysis | | 1.00 | 100.00 | 0.00 | 100.00 | 0.00 | |
| AiR Students | | 1.00 | 100.00 | 0.00 | 100.00 | 0.00 | |
| Communication between pr... | | 1.00 | 100.00 | 0.00 | 100.00 | 0.00 | |
| Learning something new | | 1.00 | 100.00 | 0.00 | 100.00 | 0.00 | |
| Mentoring experience | | 1.00 | 100.00 | 0.00 | 100.00 | 0.00 | |
| Negative | | 0.53 | 93.65 | 4.04 | 89.61 | 6.35 | |
| Negative | | 0.56 | 93.54 | 4.78 | 88.76 | 6.46 | |
| Negative | | 0.49 | 96.80 | 1.67 | 95.13 | 3.20 | |
| Negative | | 0.75 | 98.26 | 2.70 | 95.56 | 1.73 | |
| Negative | | 0.16 | 97.10 | 0.30 | 96.80 | 2.90 | |
| Negative | | 0.19 | 92.86 | 1.01 | 91.85 | 7.15 | |
| Negative | | 0.32 | 98.48 | 0.37 | 98.11 | 1.52 | |
| Negative | | 0.01 | 98.24 | 0.01 | 98.23 | 1.76 | |
| Positive | | 0.24 | 92.98 | 1.25 | 91.73 | 7.02 | |
| Positive | | 0.19 | 91.10 | 1.31 | 89.79 | 8.90 | |
| Positive | | 0.43 | 99.65 | 0.14 | 99.51 | 0.36 | |
| Positive | | 0.27 | 99.81 | 0.04 | 99.77 | 0.19 | |
| Positive | | 0.49 | 92.29 | 4.36 | 87.93 | 7.71 | |
| Positive | | 0.55 | 89.63 | 8.18 | 81.45 | 10.37 | |
| Positive | | 0.36 | 89.18 | 3.93 | 85.25 | 10.82 | |
| Positive | | 0.42 | 87.94 | 5.46 | 82.48 | 12.06 | |
| Program design and implem... | | 1.00 | 100.00 | 0.00 | 100.00 | 0.00 | |
| Program experience | | 1.00 | 100.00 | 0.00 | 100.00 | 0.00 | |
| Program outcomes | | 1.00 | 100.00 | 0.00 | 100.00 | 0.00 | |
| Science communication to t... | | 1.00 | 100.00 | 0.00 | 100.00 | 0.00 | |
| Time or length of the program | | 1.00 | 100.00 | 0.00 | 100.00 | 0.00 | |
| Virtual or remote nature of e... | | 1.00 | 100.00 | 0.00 | 100.00 | 0.00 | |
| Way of thinking | | 1.00 | 100.00 | 0.00 | 100.00 | 0.00 | |

Coding Comparison Query Criteria

Search in: **Files and Externals** **Selected Items** **Selected Folders or Static Sets** **Files with Classifications**

Coded to: All Codes

User Group A: Reviewer 3

User Group B: Reviewer 2

Unweighted Values | Weighted Values

**Overall Unweighted Kappa:** 0.42

| Name | File Size (characters) | Kappa | Agreeme... | A and B (%) | Not A and Not B (... | Disagreem... | A and N |
|---|---|---|---|---|---|---|---|
| AiR Faculty | | 1.00 | 100.00 | 0.00 | 100.00 | 0.00 | |
| AiR Program | | 1.00 | 100.00 | 0.00 | 100.00 | 0.00 | |
| AiR program analysis | | 1.00 | 100.00 | 0.00 | 100.00 | 0.00 | |
| AiR Students | | 1.00 | 100.00 | 0.00 | 100.00 | 0.00 | |
| Communication between pr... | | 1.00 | 100.00 | 0.00 | 100.00 | 0.00 | |
| Learning something new | | 1.00 | 100.00 | 0.00 | 100.00 | 0.00 | |
| Mentoring experience | | 1.00 | 100.00 | 0.00 | 100.00 | 0.00 | |
| Negative | | 0.37 | 93.49 | 2.24 | 91.25 | 6.52 | |
| Negative | | 0.53 | 94.07 | 3.84 | 90.23 | 5.94 | |
| Negative | | 0.25 | 96.48 | 0.62 | 95.86 | 3.52 | |
| Negative | | 0.68 | 97.84 | 2.42 | 95.42 | 2.16 | |
| Negative | | 0.15 | 99.34 | 0.06 | 99.28 | 0.66 | |
| Negative | | 0.38 | 96.79 | 1.05 | 95.74 | 3.22 | |
| Negative | | 0.55 | 99.07 | 0.58 | 98.49 | 0.93 | |
| Negative | | 0.09 | 99.76 | 0.01 | 99.75 | 0.24 | |
| Positive | | 0.14 | 90.61 | 0.93 | 89.68 | 9.38 | |
| Positive | | 0.15 | 90.91 | 1.03 | 89.88 | 9.09 | |
| Positive | | 0.21 | 99.50 | 0.07 | 99.43 | 0.50 | |
| Positive | | 0.30 | 99.60 | 0.09 | 99.51 | 0.41 | |
| Positive | | 0.44 | 91.66 | 3.86 | 87.80 | 8.34 | |
| Positive | | 0.48 | 86.78 | 8.28 | 78.50 | 13.23 | |
| Positive | | 0.40 | 90.12 | 4.02 | 86.10 | 9.89 | |
| Positive | | 0.36 | 87.97 | 4.36 | 83.61 | 12.03 | |
| Program design and implem... | | 1.00 | 100.00 | 0.00 | 100.00 | 0.00 | |
| Program experience | | 1.00 | 100.00 | 0.00 | 100.00 | 0.00 | |
| Program outcomes | | 1.00 | 100.00 | 0.00 | 100.00 | 0.00 | |
| Science communication to t... | | 1.00 | 100.00 | 0.00 | 100.00 | 0.00 | |
| Time or length of the program | | 1.00 | 100.00 | 0.00 | 100.00 | 0.00 | |
| Virtual or remote nature of e... | | 1.00 | 100.00 | 0.00 | 100.00 | 0.00 | |
| Way of thinking | | 1.00 | 100.00 | 0.00 | 100.00 | 0.00 | |

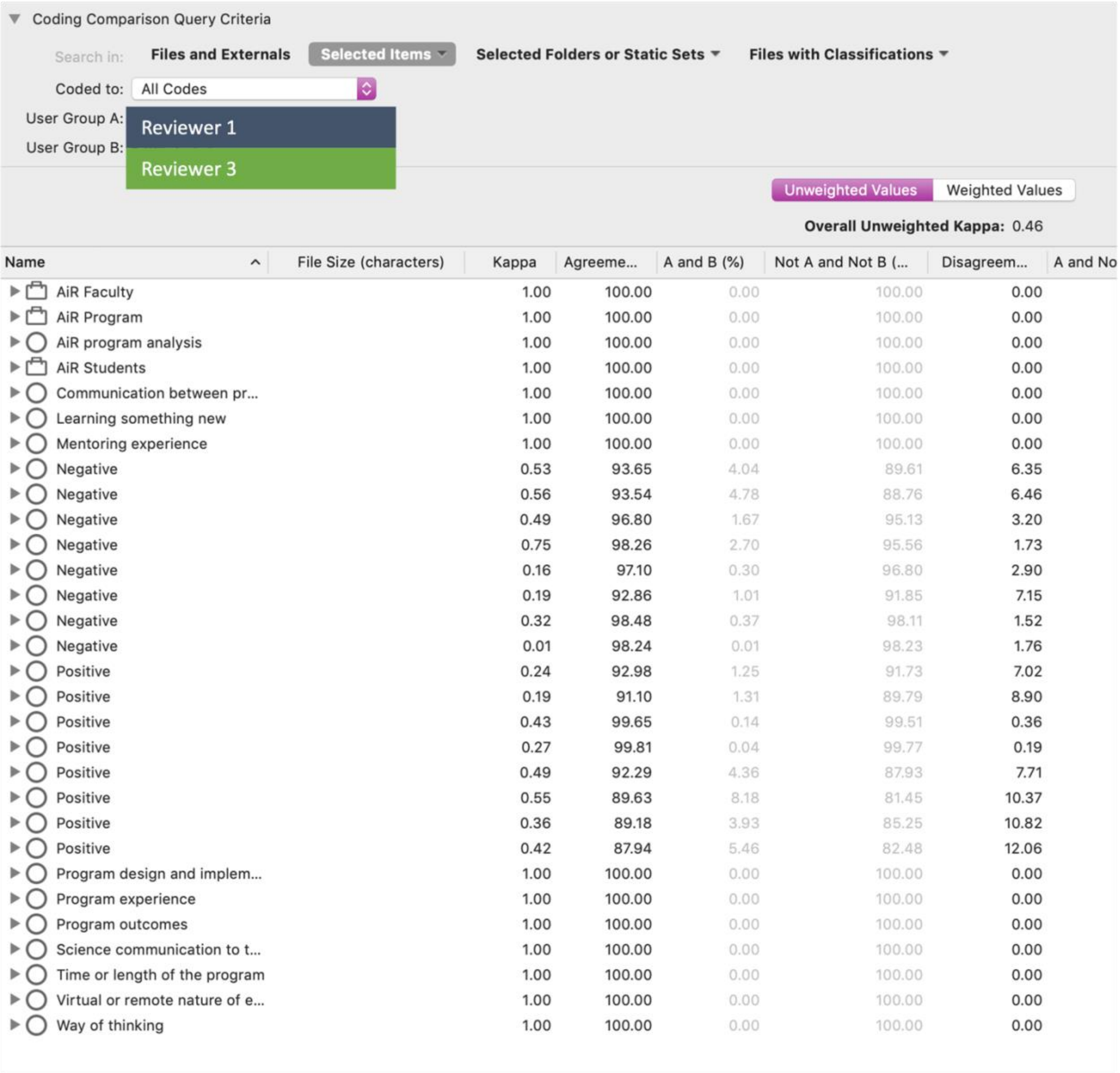


| Name | File Size (characters) | Kappa | Agreeme... | A and B (%) | Not A and Not B (... | Disagreem... | A and No |
|---|---|---|---|---|---|---|---|
| AiR Faculty | | 1.00 | 100.00 | 0.00 | 100.00 | 0.00 | |
| AiR Program | | 1.00 | 100.00 | 0.00 | 100.00 | 0.00 | |
| AiR program analysis | | 1.00 | 100.00 | 0.00 | 100.00 | 0.00 | |
| AiR Students | | 1.00 | 100.00 | 0.00 | 100.00 | 0.00 | |
| Communication between pr... | | 1.00 | 100.00 | 0.00 | 100.00 | 0.00 | |
| Learning something new | | 1.00 | 100.00 | 0.00 | 100.00 | 0.00 | |
| Mentoring experience | | 1.00 | 100.00 | 0.00 | 100.00 | 0.00 | |
| Negative | | 0.53 | 93.65 | 4.04 | 89.61 | 6.35 | |
| Negative | | 0.56 | 93.54 | 4.78 | 88.76 | 6.46 | |
| Negative | | 0.49 | 96.80 | 1.67 | 95.13 | 3.20 | |
| Negative | | 0.75 | 98.26 | 2.70 | 95.56 | 1.73 | |
| Negative | | 0.16 | 97.10 | 0.30 | 96.80 | 2.90 | |
| Negative | | 0.19 | 92.86 | 1.01 | 91.85 | 7.15 | |
| Negative | | 0.32 | 98.48 | 0.37 | 98.11 | 1.52 | |
| Negative | | 0.01 | 98.24 | 0.01 | 98.23 | 1.76 | |
| Positive | | 0.24 | 92.98 | 1.25 | 91.73 | 7.02 | |
| Positive | | 0.19 | 91.10 | 1.31 | 89.79 | 8.90 | |
| Positive | | 0.43 | 99.65 | 0.14 | 99.51 | 0.36 | |
| Positive | | 0.27 | 99.81 | 0.04 | 99.77 | 0.19 | |
| Positive | | 0.49 | 92.29 | 4.36 | 87.93 | 7.71 | |
| Positive | | 0.55 | 89.63 | 8.18 | 81.45 | 10.37 | |
| Positive | | 0.36 | 89.18 | 3.93 | 85.25 | 10.82 | |
| Positive | | 0.42 | 87.94 | 5.46 | 82.48 | 12.06 | |
| Program design and implem... | | 1.00 | 100.00 | 0.00 | 100.00 | 0.00 | |
| Program experience | | 1.00 | 100.00 | 0.00 | 100.00 | 0.00 | |
| Program outcomes | | 1.00 | 100.00 | 0.00 | 100.00 | 0.00 | |
| Science communication to t... | | 1.00 | 100.00 | 0.00 | 100.00 | 0.00 | |
| Time or length of the program | | 1.00 | 100.00 | 0.00 | 100.00 | 0.00 | |
| Virtual or remote nature of e... | | 1.00 | 100.00 | 0.00 | 100.00 | 0.00 | |
| Way of thinking | | 1.00 | 100.00 | 0.00 | 100.00 | 0.00 | |

**Supplementary Figure 4.** Program evaluation qualitative data.

Experience: 1,058 coding references

#### Frequency

| | Codes | AiR Faculty | AiR Students | Total |
|---|---|---|---|---|
| Communication | Negative | 16.2% | 15.53% | 15.69% |
| | Positive | 23.24% | 20.43% | 21.08% |
| Program design | Negative | 11.97% | 18.51% | 16.99% |
| | Positive | 26.76% | 25.53% | 25.82% |
| Time | Negative | 7.75% | 11.91% | 10.95% |
| | Positive | 1.41% | 2.34% | 2.12% |
| Virtual | Negative | 11.27% | 4.26% | 5.88% |
| | Positive | 1.41% | 1.49% | 1.47% |
| | Total | 100% | 100% | 100% |

#### Count

| | Codes | AiR Faculty | AiR Students | Total |
|---|---|---|---|---|
| Communication: 403 | Negative | 23 | 73 | 96 |
| | Positive | 33 | 96 | 129 |
| Program design: 414 | Negative | 17 | 87 | 104 |
| | Positive | 38 | 120 | 158 |
| Time: 138 | Negative | 11 | 56 | 67 |
| | Positive | 2 | 11 | 13 |
| Virtual: 103 | Negative | 16 | 20 | 36 |
| | Positive | 2 | 7 | 9 |
| | Total | 142 | 470 | 612 |

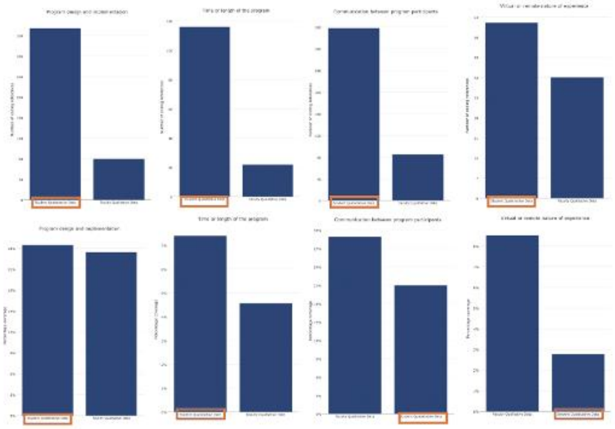

**Supplementary Figure 5.** Program outcomes qualitative data.

Outcomes: 1,685 coding reference

#### Frequency

| | Codes | AiR Faculty | AiR Students | Total |
|---|---|---|---|---|
| Learning | Negative | 16.2% | 15.53% | 15.69% |
| | Positive | 23.24% | 20.43% | 21.08% |
| Mentoring | Negative | 11.97% | 18.51% | 16.99% |
| | Positive | 26.76% | 25.53% | 25.82% |
| Science communication | Negative | 7.75% | 11.91% | 10.95% |
| | Positive | 1.41% | 2.34% | 2.12% |
| Way of thinking | Negative | 11.27% | 4.26% | 5.88% |
| | Positive | 1.41% | 1.49% | 1.47% |
| | Total | 100% | 100% | 100% |

#### Count

| | Codes | AiR Faculty | AiR Students | Total |
|---|---|---|---|---|
| Learning: 532 | Negative | 23 | 73 | 96 |
| | Positive | 33 | 96 | 129 |
| Mentoring: 490 | Negative | 17 | 87 | 104 |
| | Positive | 38 | 120 | 158 |
| Science communication: 249 | Negative | 11 | 56 | 67 |
| | Positive | 2 | 11 | 13 |
| Way of thinking: 414 | Negative | 16 | 20 | 36 |
| | Positive | 2 | 7 | 9 |
| | Total | 142 | 470 | 612 |

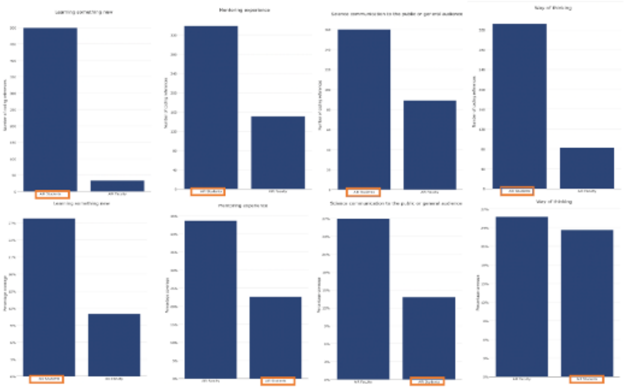
